# Parallel elevational replacement of hosts and parasites in a highly susceptible songbird genus

**DOI:** 10.1101/2025.10.10.681729

**Authors:** Daniele L. F. Wiley, Jessie L. Williamson, Silas E. Fischer, Selina M. Bauernfeind, Henry M. Streby, Kathy Granillo, Christopher C. Witt, Lisa N. Barrow

## Abstract

Elevational replacement distribution patterns underpin montane diversity and reflect the interaction of both biotic and abiotic pressures, but the degree to which parasites exhibit elevational zonation remains unclear. Investigating infection patterns in related host species across elevational gradients can reveal whether parasites and hosts show concordant patterns of elevational turnover, potentially due to shared historical and ecological factors. Here, we assessed patterns of elevational replacement in haemosporidian parasite assemblages that infect three congeneric songbird species: Bell’s vireo (*Vireo bellii),* gray vireo *(V. vicinior),* and plumbeous vireo (*V. plumbeus*), each of which breeds across distinct elevations and habitats in the southwestern United States. We screened a total of 248 individuals using cytochrome *b* PCR and microscopy. We identified 19 haemosporidian haplotypes, including eight novel lineages. We found that each of the three vireo species exhibited high haemosporidian prevalence (55.0–86.2%), with nearly all infections from the genus *Haemoproteus* (subgenus *Parahaemoproteus*). Haemosporidian assemblages varied across elevations; each sampled range of elevations harbored abundant, yet host-specific lineages with different environmental associations. Bell’s and plumbeous vireos, but not gray vireos, hosted several phylogenetically distinct, putative generalist lineages, likely reflecting spillover from more diverse local breeding bird communities. Repeated infections in individuals across breeding seasons, together with moderate parasitemia (x□ ≈ 1%) suggest that these focal vireo species harbor chronic infections during their respective breeding seasons. These results demonstrate that elevational replacement patterns in avian hosts may be mirrored by their haemosporidian parasites, particularly among host-specialized lineages.

## Introduction

Mountains harbor exceptional biodiversity, in part because elevational gradients tend to be finely partitioned by related species with ranges that are elevationally parapatric (Cadena and Loiselle, 2007; von Humboldt and Bonpland, 2009; Freeman et al., 2016). Yet, it remains unclear whether parasites of elevational replacement host species mirror this range stratification. Generally, host-parasite interactions can strongly influence biogeographic patterns, with reciprocal effects on fitness (Combes, 1997, 2001; Poulin, 2011; Hasik and Siepielski, 2022), population dynamics (Albon et al., 2002), and community structure (Minchella and Scott, 1991, McNew et al., 2021).

Strong host-specificity, combined with shared environmental constraints, should result in parasite turnover that parallels the turnover of hosts along environmental gradients. Conversely, aspects of parasite life-history––such as insulation from climatic factors (e.g., endoparasites), dependency on secondary hosts and vectors (e.g., complex life cycle), and degree of host-specificity (e.g., host range and degree of host switching)––should be expected to cause discordant patterns from those observed in hosts (Gage et al. 2008; Krasnov and Poulin, 2010; Ellis et al., 2015).

Regional surveys across elevational gradients reflect these complex relationships. In various mountain systems, parasite turnover has been found to exceed host turnover (Galen and Witt, 2014; Barrow et al., 2021), and host turnover has been found to predict parasite turnover (Ellis et al., 2015; Williamson et al., 2019). Gradients of precipitation and temperature, as well as landscape features, have also been found to shape parasite communities (Illera et al., 2017; McNew et al., 2021).

Investigating the parasite communities of closely related host taxa that are elevational replacements provides a natural experiment for disentangling the relative roles of host associations, parasite traits, and environmental gradients in shaping parasite biogeography, while controlling for host phylogeny, life history, and parasite susceptibility (Medeiros et al., 2013; Barrow et al., 2019).

Among blood parasites, haemosporidians (order Haemosporida; Danilewsky, 1885) are a cosmopolitan group of protozoans with complex life cycles dependent upon vertebrate taxa, including birds, mammals, and reptiles (Ricklefs and Fallon, 2002; Duval et al., 2007; Boysen et al., 2022), and invertebrate vectors in the order Diptera (Linnaeus, 1758). A long history of co-evolution with avian hosts (Ricklefs et al., 2004; Lauron et al., 2015; Galen et al., 2018) has given rise to a remarkable diversity of avian haemosporidian lineages (Valkiūnas, 2005; Hellgren et al., 2007; Perkins, 2014), with many still yet to be described (Borner et al., 2016). This diversity is reflected both phylogenetically and phenotypically, with variable degrees of pathogenicity (reviewed in Bennett et al., 1993) and host specificity (Ellis and Bensch, 2018) recorded among and within genera.

Given the high diversity and complexity of haemosporidian parasites, it has remained challenging to characterize the distribution patterns of haemosporidian lineages across large geographic scales (Clark, 2018). However, within regions and genera, spatial trends appear—reflecting the influence of climate directly, through physiological interactions during transmission and development (Ikemoto 2008; Mordecai et al., 2013), and indirectly, by influencing host and vector abundance and susceptibility (Filion et al., 2020). As a result, the fluctuating climate of temperate and montane regions produce seasonal, annual, and habitat-level waves of infection (Bensch et al., 2007; Pérez-Rodríguez et al., 2013; Lutz et al., 2015; Reinoso-Pérez et al., 2024), whereas areas with higher, less seasonal temperatures, such as tropical regions and lowlands, can have persistent haemosporidian pressure (Zamora-Vilchis et al., 2012).

Infection intensity, commonly measured as parasitemia (defined as the ‘estimated percentage of haemosporidians circulating in host blood’) provides insight into host condition and stage of haemosporidian infection (i.e., acute or chronic; Valkiūnas, 2005). Like prevalence (here defined as ‘proneness to infection’), parasitemia is also influenced by environmental factors, with peaks in temperate regions during the breeding season, when warmer, wetter conditions favor vector activity (Valkiūnas, 2005; Reinoso-Pérez et al., 2024) and when chronically infected hosts experience tissue-to-blood recrudescence (Atkinson and van Riper, 1991). Therefore, montane regions provide a valuable system for studying avian haemosporidian infection dynamics because of the sharp local variation in both climate and habitat over relatively small geographic distance (e.g., Illera et al., 2017; Rodríguez-Hernández et al., 2021; Williamson et al., 2019).

The southwestern U.S. presents a particularly compelling region for studying haemosporidian biogeography and dynamics due to its climatic heterogeneity, wide range of habitat types, and steep elevational gradients in sky island mountains. Although arid overall, the region exhibits strong variation in precipitation timing and intensity, shaped by both temperate seasonality and the North American monsoon system. These climatic patterns interact with topographic variation (reviewed in Coblentz and Riitters, 2004) to create a mosaic of ecologically distinct elevational zones ranging from lowland riparian corridors with high relative humidity and permanent water sources, to mid-elevation desert scrublands and savannas reliant on seasonal rainfall, to cooler montane forests with shorter growing seasons and higher annual precipitation (Sheppard et al., 2002). While community-level surveys suggest moderate overall prevalence of haemosporidian infection among avian communities in the region (mean: ∼36%; Barrow et al., 2021), and while environmental conditions––particularly related to elevation, have been shown to strongly influence spatial patterns of infection (Williamson et al., 2019)––previous studies have primarily described environmental and ecological contributions across mid-high elevation habitats and sky islands. Therefore, further work is needed to expand our understanding of whether observed environmental and elevational trends persist across broader gradients and habitat types, especially among particularly susceptible hosts.

Among the many avian species previously surveyed in the desert Southwest, the plumbeous vireo (*Vireo plumbeus*) has consistently stood out for its high haemosporidian prevalence across sites and years (Marroquin-Flores et al., 2017; Barrow et al., 2021). Migratory populations of this insectivorous songbird breed in mid- to high-elevation coniferous woodlands (from ∼1,200–3,000 m) across the western U.S. and Mexico (Barlow, 1977; Curson and Goguen, 1998) and spend the nonbreeding season along the Pacific Slope and in the lowlands of central Mexico (Sibley and Monroe, 1990). Two closely related migratory vireos, Bell’s vireo (*Vireo bellii*) and gray vireo (*Vireo vicinior*), have relatively similar breeding distributions, yet inhabit distinct habitats and elevations. gray vireos breed in arid mid-elevation juniper (*Juniperus* spp.) savannas and chaparral, ∼400–1,900 m (Hubbard, 1970; Barlow, 1977) and spend the nonbreeding season primarily in lowland desert and coastal areas of Sonora and Baja California (Barlow et al., 1999; Fischer et al., 2025). Bell’s vireos breed in lowland riparian habitats, generally below ∼1,500 m, with comparatively high access to water and vegetative cover (Brown, 1993) and spend the nonbreeding season along the Pacific coast of Mexico, from Baja California and Sonora south to El Salvador.

Given the similarities in ecology and life history among vireos, local differences in breeding elevational range make these three focal species an ideal comparative system through which to examine haemosporidian infection patterns across elevational gradients.

In this study, we tested whether haemosporidian biogeography, community assemblage, and infection dynamics vary predictably among three species of vireos that occupy distinct elevations and habitats at sites in New Mexico and Utah, USA. Using a combination of molecular screening and microscopy, we (1) compared parasite prevalence and infection intensity among host species, including the first parasite survey for regionally vulnerable gray vireos; (2) characterized haemosporidian haplotypes and host specificity; and (3) tested the extent to which community composition was linked to elevational zones and/or their associated host species and habitats. We predicted finding extensive parasite sharing between elevational replacement vireo species, considering the proximity and interdigitated nature of their ranges. Based on previously published haemosporidian surveys in vireos, we anticipated finding high infection prevalence across all three host species; however, we also expected prevalence and parasitemia to vary among species and environments. Lastly, because the Southwest is arid, we predicted higher prevalence and parasite diversity in more mesic habitats suitable for dipterid vectors, such as in the low-elevation riparian corridors used by Bell’s vireos and high-elevation woodlands occupied by plumbeous vireos.

## Materials and methods

### Sample collection

We sampled 248 wild-caught individuals of three vireo species (Bell’s vireo: *n* = 20, gray vireo: *n* = 170, and plumbeous vireo: *n* = 58; Fig. 1A; Table S1). Sampled individuals included adults and juveniles (Table S1). All individuals were sampled during the summer months (May–August), with the majority sampled prior to monsoon season (∼early-July–September) from 1995–2019 across 28 unique sites in New Mexico and a single site in Utah. During 2017–2019, targeted fieldwork was conducted to sample gray vireos: most individuals were captured in mist nets, with the exception of hatch-year birds sampled at nests. Blood samples were taken from the brachial vein and stored in lysis buffer, and individuals were released at capture sites for ongoing study of gray vireo annual cycle ecology (Fischer, 2020; Fischer et al., 2022; Fischer et al., 2025). In 2018 only, we prepared and air-dried blood smears in the field (*n* = 53). We also recaptured and collected a second whole blood sample from fourteen gray vireos (approximately one year after their initial sample collection in 2017), yielding two total whole blood samples and one smear for this subset of individuals (Table S2). Additional gray vireo tissues for four individuals were deposited at the Museum of Southwestern Biology (hereafter, MSB) at the University of New Mexico following protocols approved by the UNM Institutional Animal Care and Use Committee. Samples for Bell’s and plumbeous vireos were requested as tissue loans from the MSB. Detailed sampling protocol for Bell’s and plumbeous vireos are described in (Marroquin-Flores et al., 2017; Barrow et al., 2021; Gyllenhaal, 2024).

**Figure 1.**
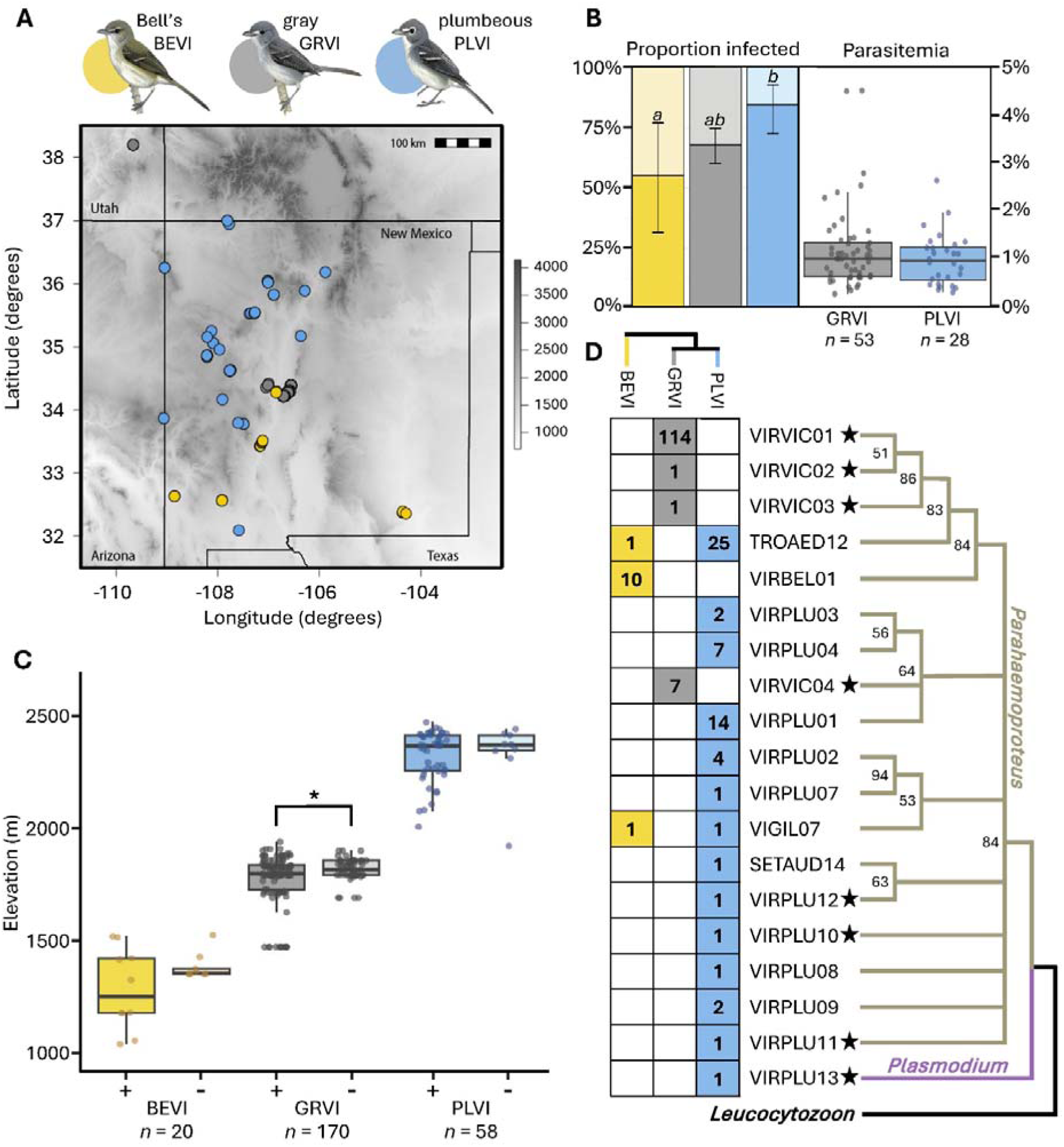
A) Sampling distribution for each host species (Bell’s vireo: *Vireo bellii,* gray vireo: *V. vicinior,* and plumbeous vireo: *V. plumbeus*) in the southwestern USA; B) Left: summary of infection prevalence, with significant differences between groups indicated by letters. Error bars represent 95% confidence intervals for positive infection proportions. Right: percent parasitemia for gray and plumbeous vireos. C) Proportion of individuals infected versus uninfected across elevational gradient. Asterisk denotes significant p-values (< 0.05); D) Phylogeny of haemosporidian cyt *b* haplotypes and their respective abundances by host. Stars denote novel haplotypes and bootstrap values >50 are shown at nodes. Bird images courtesy Birds of the World (Billerman et al., 2025).

All sampled birds either had specific coordinates recorded at the time of capture or were caught within 100 m of an existing sampling location. For the latter, we assigned geographically proximate coordinates based on the nearest documented sampling site or, when birds were captured between two closely spaced sites, by taking the midpoint between them (Table S3).

### Parasite screening and genetic data collection

We extracted DNA from pectoral muscle (*n =* 82) and whole blood (*n =* 166) using the QIAGEN DNeasy Blood and Tissue Kit following manufacturer’s recommendations. Birds were screened for *Haemoproteus* and *Plasmodium* parasites using three nested polymerase chain reaction (PCR) protocols to amplify a 478-base pair (bp) fragment of the haemosporidian mitochondrial cyt *b* gene (Hellgren et al., 2004; Waldenström et al., 2004). We amplified parasite DNA with the outer primer pairs HaemNFI/HaemNR3 and HaemNF/HaemNR2, followed by the nested primer pair HaemF/HaemR2. We prepared outer PCR reactions in 25 μL volumes, containing 1.25 U of AmpliTaq Gold DNA Polymerase (Applied Biosystems, Foster City, CA, USA), 1× PCR Buffer II, 2.5 mM MgCl2, 0.2 mM dNTPs, 0.5 μM of each primer, and 20 ng of template DNA. Thermal cycling conditions were modified from Galen and Witt (2014), with an initial denaturation at 95 °C for 8 mins, followed by 20 cycles of 94 °C for 30 seconds, 50 °C for 30 secs, 72 °C for 45 secs, and a final extension at 72 °C for 10 mins.

For nested PCR, we used 1 μL of the outer PCR product as the template, with reaction conditions identical to the outer PCR, except for an increase to 35 cycles. Each reaction set included negative and positive controls to monitor for contamination and confirm amplification success, respectively. We visualized PCR products on 2% agarose gels stained with SYBR Safe Gel Stain (Invitrogen, Carlsbad, CA, USA) to verify the presence of amplicons of the expected size.

Successfully amplified products were purified using ExoSap-IT (Affymetrix, Inc., Santa Clara, CA, USA) and submitted for Sanger sequencing at Psomagen (Rockville, MD, USA). gray vireo blood samples from recaptured individuals (i.e., samples from the same individuals from 2017 and 2018) underwent PCR amplification, but only a single sample was sequenced (Table S2).

### Determining prevalence and haplotype diversity

We determined positive infections upon successful amplification of haemosporidian cyt *b* sequences and calculated pathogen prevalence within each host species and calculated 95% binomial confidence intervals using the ‘exact’ method available in the *binom* package in R (version 4.3.2; R Core Team, 2023; Sundar, 2006).

Haemosporidian cyt *b* forward and reverse reads were trimmed to remove primers, resulting in the target fragment size of 478 bp and were then assembled using the default alignment algorithm in Geneious (version 2025.03; Biomatters Ltd; Kearse et al., 2012; Table S4). To identify haplotypes, we compared cleaned sequences to published records stored in the public databases GenBank (National Center for Biotechnology Information, US National Library of Medicine) and MalAvi (Bensch et al., 2009) by using the Basic Local Alignment Search Tool (BLAST).

Additionally, to ensure the accuracy of both established and novel haplotypes, we downloaded all haemosporidian cyt *b* haplotype sequence files from MalAvi and compared the number of differences, if any, to our sequences via a distance matrix calculated in Geneious. Haplotypes that differed by one or more base pairs (∼0.2% sequence divergence) from published sequences on GenBank or MalAvi were considered novel and named following MalAvi conventions.

### Phylogenetic analysis

We estimated phylogenetic relationships among haemosporidian haplotypes using a maximum-likelihood (ML) framework in RAxML, v8.2.10 (Stamatakis, 2014). Using the GTR+G model of nucleotide substitution, we conducted a rapid bootstrap analysis with 1000 replicates, after which we searched for the best-scoring ML tree. We rooted the tree with *Leucocytozoon* (COLBF21, GenBank Accession MK947795) based on the current phylogenetic hypothesis for avian haemosporidians (Borner et al., 2016; Galen et al., 2018). Nodes with <50% bootstrap support were collapsed in TreeGraph2 (Stöver and Müller, 2010). We determined the relationships among the three vireo study species based on the posterior distribution of likely trees available on birdtree.org with the ‘Hackett All Species’ option (Hackett et al., 2008; Jetz et al., 2012). We downloaded 100 phylogeny subsets and selected the first tree because the relationships were consistent among the 100 trees.

### Estimating parasite species richness and sampling completeness

We used the *iNEXT* R package (version 3.0.1; Chao et al., 2014; Hsieh et al., 2024) to estimate parasite lineage diversity and evaluate sampling completeness for host species with adequate data (i.e., gray and plumbeous vireos). In separate analyses for each host species, we used infection counts of each parasite haplotype to generate rarefaction and extrapolation curves for parasite species richness (q = 0), with the endpoint set at 400 individuals (i.e., infections). We applied the ‘iNEXT()’ function to compute species richness estimates, including observed richness, extrapolated richness, and associated confidence intervals using standard function parameters.

Additionally, we used the output to estimate sampling completeness (Chao et al., 2014) and defined this value as the minimum number of individual birds required to detect 95% of the total estimated parasite haplotypes infecting each host species. To visualize the rarefaction curves, we used the ‘ggiNEXT()’ function and *ggplot2* package for style modifications (Wickham, 2016).

### Microscopy and parasitemia calculations

We obtained available blood smears from the MSB for plumbeous vireos (*n =* 28) and made and air dried blood smears in the field for gray vireos (*n =* 53). Each slide was fixed with absolute methanol and stained for 50 mins with Giemsa solution (pH 7.0; Sigma-Aldrich, St. Louis, MO, USA). We then examined each slide to confirm infection status using light microscopy on an Olympus BX 53 Microscope following the protocol described in Valkiūnas (2005), where 10,000 erythrocytes were scanned at 1000x magnification with an oil immersion lens to identify and count *Parahaemoproteus* and *Plasmodium* infections (Valkiūnas, 2005). We took digital photographs of representative haemosporidian specimens, which were archived at the MSB and are searchable via the Arctos database (https://arctos.database.museum/). We calculated parasitemia, or the estimated percentage of erythrocytes infected with haemosporidian parasites, as the number of infected erythrocytes out of 10,000 screened. To account for lack of normality in the data, we used a bootstrap method with 1,000 replications to calculate parasitemia 95% confidence intervals via the R package *boot* (Kushary, 2000).

### Characterizing environmental variation

We characterized elevation (m) and climatic variation using 19 bioclimatic variables at 30 sec (∼1 km²) resolution from the WorldClim 2.1 database (Fick and Hijmans, 2017). We used principal components analysis (PCA) for temperature (Bio1–11) and precipitation (Bio12–19) variables to create a composite measure of temperature and precipitation across the gradient. The first two axes for temperature (hereafter, tempPC1 and tempPC2) explained a combined 84.2% of the variation and the first two axes for precipitation (hereafter, precipPC1 and precipPC2) explained 95.1% of the variation (Fig. S1; Table S5-6). Variable loadings indicated that tempPC1 primarily represented year-round temperatures, with higher values relating to lower average temperatures, while tempPC2 represented greater seasonality and less temperature stability, with higher values relating to more extreme temperature fluctuations throughout the year and day-night cycle.

Similarly, precipPC1 described the overall precipitation amount, with higher values indicating wetter conditions, whereas precipPC2 represented precipitation seasonality, with higher values representing more stable and less seasonal precipitation (Table S5-6).

### Assessing infection dynamics among host species and infection types

We tested whether haemosporidian infection status differed between categorical variables such as host species and tissue type (i.e., pectoral muscle or whole blood samples), using Pearson’s chi-squared tests via the *stats* package. To examine the relative magnitude of effect for each comparison, we calculated Cramér’s V effect sizes were using the ‘CramerV()’ function in the *rcompanion* package (Mangiafico, 2024). Next, to assess differences in parasitemia, we first normalized the data using a natural log ln(1+x) transformation and removed two influential outliers with parasitemia >4% (gray vireo IDs: 272132427 and 272132642). We then evaluated residual variance with the ‘var.test()’ function in the *stats* package and compared log-transformed parasitemia between gray and plumbeous vireos and infection types (i.e., single vs. coinfected) using two-tailed Student’s *t*-tests.

### Intraspecific infection prevalence and parasitemia across environments

We employed a two-pronged approach to assess environmental associations with infection prevalence: First, we used univariate, non-parametric tests to characterize and compare intraspecific infection patterns across abiotic variables relevant to parasite and vector ecology, including elevation, overall temperature (tempPC1) and precipitation (precipPC1), and the seasonality of temperature (tempPC2) and precipitation (precipPC2). Then, we built intraspecific multivariate linear models and applied nested and exhaustive model selection to evaluate the strength and direction of environmental effects. We adopted this approach to account for the structure of our data (i.e., non-overlapping elevational and environmental bands distinctly correlated to species identity) and limitations of our dataset (i.e., limited sample sizes, degree of environmental variation, and infection class imbalances).

We first assessed data normality both visually and then statistically, using the Shapiro-Wilk test using the ‘shapiro.test()’ function from the *stats* package. Given the non-normal distribution of environmental and geographic data in our dataset, we applied a rank-sum approach via Mann-Whitney U-tests to compare the distributions of environmental variables between infected and uninfected birds. Wilcoxon effect sizes of each comparison were then calculated via the ‘wilcox_effsize()’ function in the *rstatix* package (Kassambara, 2023).

We then constructed species-specific generalized linear models using glm() from the *stats* package. Prevalence model sets used a binomial distribution (hereafter: “BEVI_prev” for Bell’s vireo, “GRVI_prev” for gray vireo, and “PLVI_prev” for plumbeous vireo sets), while parasitemia model sets used a Gaussian distribution (“GRVI_para”, “PLVI_para”). We evaluated linearity between the logit and predicted values in prevalence models and assessed residual diagnostics in parasitemia models and with ‘autoplot()’ within the *ggfortify* package (Tang et al., 2016).

Multicollinearity between variables was assessed by calculating variance inflation factors (VIF) via the *car* package. As expected, there was high collinearity with aspects of geographic and topographic variation, i.e. latitude and elevation, with other environmental variables. We retained elevation in all model sets because it captured variation in habitat and vector dynamics important to infection that were not explained by temperature and precipitation alone (Ishtiaq and Barve, 2018); however, we removed latitude from all linear models to reduce overfitting.

For model sets with adequate sampling (i.e., GRVI_prev, PLVI_prev, GRVI_para, PLVI_para), we evaluated all possible additive combinations of five predictor variables: elevation, tempPC1, tempPC2, precipPC1, precipPC2. We used the ‘dredge’ function in the *MuMIn* package (Bartoń, 2024) for exhaustive model selection. For host datasets with < 30 data points (BEVI_prev and PLVI_para) we restricted candidate models to simple, biologically relevant structures to avoid overfitting. Specifically, we included univariate models nested within simple additive models containing at most two predictor variables, as well as null and global models for comparison. We then compared model performance and evaluated the trade-off between model fit and complexity using Akaike’s Information Criterion corrected for small sample sizes (AICc; Table 1).

**Table 1.** Top five models assessing environmental predictors of haemosporidian infection for each species. Sample sizes for datasets and directionality of best model outputs are listed at the top. Models were ranked by AICc, with ΔAICc values relative to the top model and corresponding Akaike weights (Weight) indicating model support.

| *Top five infection probability fixed effects additive models* |  |  |  |  |  |
| --- | --- | --- | --- | --- | --- |
| **Bell's vireo (*n* = 20)** |  |  |  |  |  |
| *Rank* | *Predictors* | *K* | *AICc* | *ΔAICc* | *Weight* |
| 1 | tempPC2 + precipPC1 | 3 | 22.6 | - | 0.324 |
| 2 | tempPC1 | 2 | 23.6 | 0.967 | 0.200 |
| 3 | precipPC1 | 2 | 24.7 | 0.348 | 0.113 |
| 4 | tempPC1 + precipPC1 | 3 | 25.9 | 0.199 | 0.064 |
| 5 | Elevation + tempPC1 | 3 | 25.9 | 0.197 | 0.064 |
| **gray vireo (*n* = 170)** |  |  |  |  |  |
| *Rank* | *Predictors* | *K* | *AICc* | *ΔAICc* | *Weight* |
| 1 | Elevation + tempPC1 | 3 | 204 | - | 0.419 |
| 2 | Elevation + precipPC1 + tempPC1 | 4 | 206 | 1.72 | 0.177 |
| 3 | Elevation + precipPC2 + tempPC1 | 4 | 206 | 1.73 | 0.177 |
| 4 | Elevation + tempPC1 + tempPC2 | 4 | 206 | 2.07 | 0.149 |
| 5 | Elevation + tempPC1 + tempPC2, precipPC1 + precipPC2 | 5 | 207 | 3.35 | 0.078 |
| **plumbeous vireo (*n* = 58)** |  |  |  |  |  |
| *Rank* | *Predictors* | *K* | *AICc* | *ΔAICc* | *Weight* |
| 1 | Null model | 1 | 52.1 | - | 0.301 |
| 2 | precipPC2 | 2 | 52.4 | 0.409 | 0.246 |
| 3 | precipPC1 | 2 | 53.3 | 1.17 | 0.168 |
| 4 | tempPC2 | 2 | 53.4 | 1.29 | 0.158 |
| 5 | PrecipPC1 + precipPC2 | 2 | 53.9 | 1.73 | 0.127 |

<sup>2</sup> **Table 1.** Top five models assessing environmental predictors of haemosporidian infection for each species. Sample sizes for datasets and directionality of best model outputs are listed at the top. Models were ranked by AICc, with ΔAICc values relative to the top model and corresponding Akaike weights (Weight) indicating model support.

To examine the effect of each predictor in top-ranked models, we standardized and calculated model-averaged regression coefficients using the *MuMIN* package (Table 2). For top-ranked models where the null model was not favored, we assessed goodness of fit via Hosmer-Lemeshow tests as part of the *ResourceSelection* package (Lele et al., 2023) and checked for overdispersion by dividing residual deviance by the degrees of freedom. Predictive accuracy was also assessed using 10-fold cross-validation via the *caret* package (Kuhn, 2008).

**Table 2.**
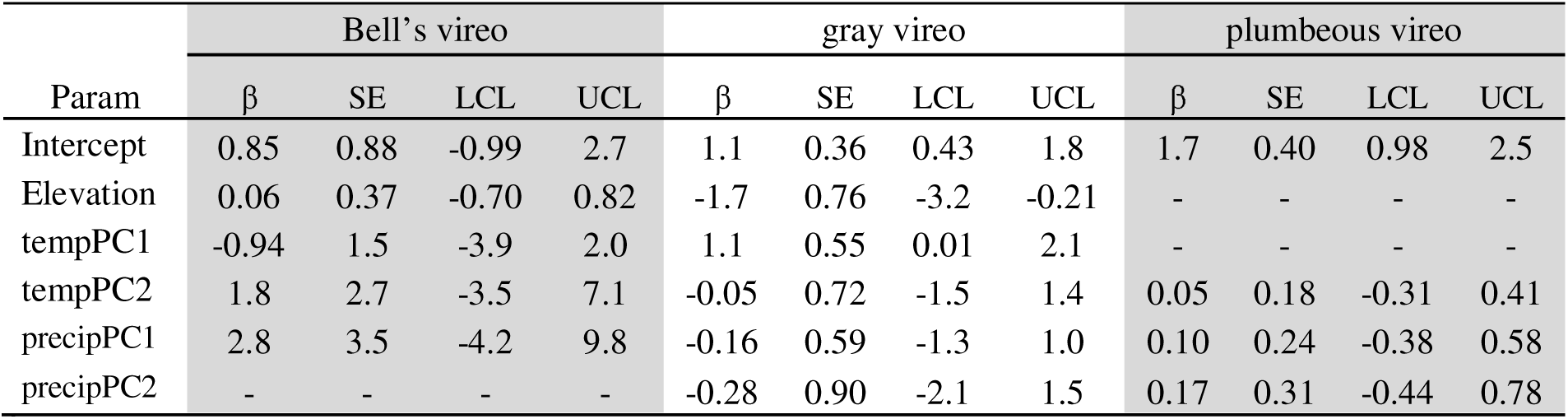
Standardized model-averaged regression coefficients used to estimate effects of predictors and precision of effects across species-specific candidate model sets that assess environmental predictors of haemosporidian infection. Regression coefficients (β), standard error (SE), and 95% confidence limits (lower as LCL, upper as UCL) are given. Dashes indicate that the parameter was not present in a final model set, and NA values indicate that the parameter was not tested due to lack of variation present in model set.

| Param | Bell's vireo |  |  |  | gray vireo |  |  |  | plumbeous vireo |  |  |  |
| --- | --- | --- | --- | --- | --- | --- | --- | --- | --- | --- | --- | --- |
| | $\beta$ | SE | LCL | UCL | $\beta$ | SE | LCL | UCL | $\beta$ | SE | LCL | UCL |
| Intercept | 0.85 | 0.88 | -0.99 | 2.7 | 1.1 | 0.36 | 0.43 | 1.8 | 1.7 | 0.40 | 0.98 | 2.5 |
| Elevation | 0.06 | 0.37 | -0.70 | 0.82 | -1.7 | 0.76 | -3.2 | -0.21 | - | - | - | - |
| tempPC1 | -0.94 | 1.5 | -3.9 | 2.0 | 1.1 | 0.55 | 0.01 | 2.1 | - | - | - | - |
| tempPC2 | 1.8 | 2.7 | -3.5 | 7.1 | -0.05 | 0.72 | -1.5 | 1.4 | 0.05 | 0.18 | -0.31 | 0.41 |
| precipPC1 | 2.8 | 3.5 | -4.2 | 9.8 | -0.16 | 0.59 | -1.3 | 1.0 | 0.10 | 0.24 | -0.38 | 0.58 |
| precipPC2 | - | - | - | - | -0.28 | 0.90 | -2.1 | 1.5 | 0.17 | 0.31 | -0.44 | 0.78 |

<sup>3</sup>
<sup>3</sup> **Table 2.** Standardized model-averaged regression coefficients used to estimate effects of predictors and precision of effects across species-specific candidate model sets that assess environmental predictors of haemosporidian infection. Regression coefficients ( $\beta$ ), standard error (SE), and 95% confidence limits (lower as LCL, upper as UCL) are given. Dashes indicate that the parameter was not present in a final model set, and NA values indicate that the parameter was not tested due to lack of variation present in model set.

## Results

### Haemosporidian prevalence, abundance, and diversity

We uncovered an unusually high prevalence of haemosporidian infections in all three vireo species, with 175 infected birds of 248 screened (70.6%; 95% CI [64.5 – 76.2%]). The highest documented prevalence was found in plumbeous vireos (84.5% infected; 95% CI [72.6 – 92.7%]), followed by gray vireos (67.7%; 95% CI [60.0 – 74.7%]), and lastly, Bell’s vireos (55%; 95% CI [31.5 – 76.9%]; Fig. 1B). Prevalence was significantly lower in Bell’s than in plumbeous vireos (χ²(1) = 5.72, p = 0.017, φ = 0.31), indicating a moderate effect size. While most birds were infected by single haemosporidian haplotype infections (88.1%; 95% CI [82.3 – 92.5%]), we found 21 individuals that were coinfected with two or more haplotypes (12.0%; 95% CI [7.54 – 17.7%]), and one gray vireo that was infected by three haplotypes (VIRVIC01, VIRVIC02, VIRVIC03). Of the 14 gray vireos recaptured about a year after initial sampling, 12 were infected with haemosporidians in both years, while two that were uninfected in 2017 tested positive in 2018.

Nearly all infections were caused by parasites in the genus *Haemoproteus*, specifically the subgenus *Parahaemoproteus* (99.5%), which comprised 18 of 19 haplotypes identified. *Plasmodium* was rare in our data, with a single infection (lineage VIRPLU13) recorded from a plumbeous vireo (MSB:Bird:60726; Table S4). Of the 19 total parasite haplotypes identified, 8 were novel (Fig. 1D; Table S4).

### Estimates of parasite species richness and sampling completeness

We identified a total of three haemosporidian haplotypes from Bell’s vireos, four from gray vireos, and 14 from plumbeous vireos (Fig. 1D). These differences were partly, but not entirely, attributable to differences in sample size. gray vireo *Parahaemoproteus* infections were substantially less diverse than those of plumbeous vireos despite more intensive screening efforts. Extrapolation from rarefaction curves showed considerably lower estimated parasite richness for gray vireos (4.99 ± 0.53 SE; 95% CI [4.00–6.03]; Table S7) and estimated that much smaller sample sizes would be needed to reach 95% sampling completeness (14 infections; Fig. 2; Table S8). This result contrasted our results for plumbeous vireos, where parasite richness was estimated to be ∼10x higher than gray vireos (53.8 ± 23.3 SE; 95% CI [14.0–99.6]) and require much greater sampling effort (∼26x more than estimated for gray vireos) to reach an estimated 95% completeness (estimated 364 infections). Bell’s vireos also exhibited low lineage diversity (3.92 ± 0.784 SE; 95% CI [3.00-5.45] and an estimated 95% sample completeness at only 33 infections, consistent with the limited diversity recovered (Table S7-S8). However, we caution overinterpreting these results due to the small Bell’s vireo sample size (*n* = 12 infected) in our study and recommend additional sampling to generate more precise estimates.

**Figure 2.**
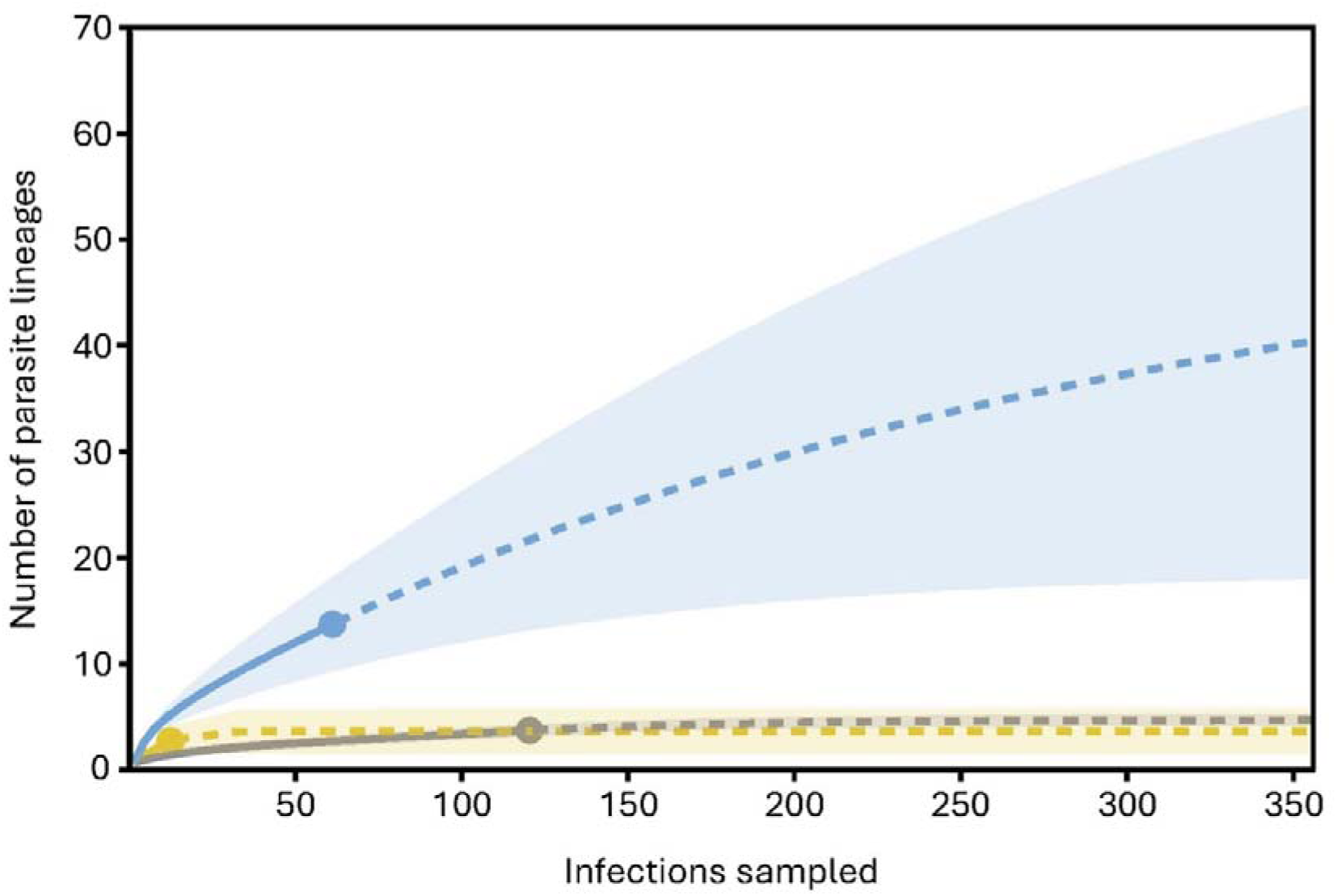
Estimate of haemosporidian lineage diversity in Bell’s (*Vireo belli*), gray (*V. vicinior*) and plumbeous vireos (*V. plumbeus*) based on iNEXT rarefaction and extrapolation. Bell’s vireo curves (yellow) were created using 12 avian haemosporidian infections and three haplotypes; gray vireo curves (gray) used 123 avian haemosporidian infections and four haplotypes; plumbeous vireo curves (blue) used 62 haemosporidian infections and 14 haplotypes. Points indicate the reference sample, solid line the rarefaction estimate, and dotted line the extrapolation. The analysis suggested that sampling approximately 33 infections in Bell’s, 14 in grays, and 364 total infections for plumbeous vireos would capture >95% of the haemosporidian lineage diversity in this community. Total haplotype richness, rounded to nearest whole number, is estimated to be four for Bell’s vireos (95% CI [3-6]), five for gray vireos (95% CI [4 – 6]) and 54 for plumbeous vireos (95% CI [14 – 101]).

Of the 19 total haplotypes identified, we found evidence of potential host specificity with at least one species-specific haplotype infecting each of the three vireo species (Table S4). Of note, gray vireos were infected by novel haemosporidian haplotypes only, (VIRVIC01-VIRVIC04). This finding contrasted with plumbeous vireos, which showed novel haplotypes (i.e. VIRPLU10-13) and haplotypes previously found only in plumbeous vireos (i.e. VIRPLU03, VIRPLU07-09), as well as lineages known to infect other non-vireo species (i.e. SETAUD14, TROAED12, VIGIL07, VIRPLU01, and VIRPLU04). Sampling was comparatively lower for Bell’s vireos; no novel lineages were detected. We did detect, however, a recently described lineage (VIRBEL01) which was previously documented in Bell’s vireos sampled from Arizona and Texas (Fecchio et al., 2023), as well as other more generalist lineages (i.e. TROAED12 and VIGIL07; Table S4).

### Intensity of infection (parasitemia) across hosts and haplotypes

We screened 102 individuals via microscopy (gray *n* = 71, plumbeous *n* = 31). A total of 81 samples showed evidence of infection and had parasitemia *calculated (Parahaemoproteus:* gray *n* = 53 and plumbeous *n* = 27; *Plasmodium*: plumbeous *n* = 1). gray vireos typically had slightly higher infection loads (1.00%; 95% CI [0.782 – 1.27%]) compared to plumbeous vireos (0.786%; 95% CI [0.584 – 1.01%]), partly due to two outlier gray vireo samples with >4% parasitemia (ID 272132427 and 27213264; Fig 1C). After removing these outliers, we found that mean parasitemia did not differ significantly between host species (t(77) = 0.656, p = 0.514). When we compared single-haplotypes and coinfections, we also found no significant difference in parasitemia between species (t(73) = -0.728, p = 0.469), though we note the sampling pool of coinfected birds was small compared to single infections (*n* = 7 and *n* = 68, respectively).

Our comparison of infections using PCR and microscopy was largely consistent, with few detection errors identified between the two methods. We detected four positively infected birds via microscopy that did not successfully amplify by PCR (Table S1). These samples had a range of parasitemia spanning 0.70% to 2.68%. Additionally, we failed to detect haemosporidian infections via microscopy for two birds that were later successfully amplified using PCR; one individual was the single coinfected gray vireo with three *Parahaemoproteous* haplotypes that were identified by sequencing (Table S1).

### Environmental patterns of infection status and parasitemia

*Parahaemoproteus* prevalence showed idiosyncratic patterns among host species and their associated habitats (Fig 3; Table S9). For example, we found that infected Bell’s vireos tended to be found at lower latitudes (*z* = -2.09, *p* = 0.040), with warmer year-round temperatures (tempPC1; *z* = -2.25, *p* = 0.027 Fig. 3; Fig. S2; Table S9). Similarly, probability of infection increased as overall precipitation (precipPC1) and temperature seasonality (tempPC2) increased (Table 1; Table 2). In contrast, we found higher proportions of infected gray vireos at lower elevations (*z* = -2.07, *p* = 0.038), Fig. 1C; Table S9) and at sites with more stable precipitation (precipPC2; z = 1.72, p = 0.057), Fig 3; Table S9). However, this result was dampened when we excluded the eleven infected gray vireos from the Abajo Mountain site in Utah, which were outliers in many of the environmental PCA dimensions. The top model, however, showed that infection was negatively associated with elevation and positively associated with temperature (tempPC1; Table 1; Table 2) even when excluding the Utah site. Lastly, plumbeous vireo infections tended to be higher in areas with more seasonally stable precipitation (precipPC2; *z* = 2.04, *p* = 0.042), Fig. 3). Despite the top-ranked infection probability model showing no significant linear trends, precipitation (precipPC1 and precipPC2) was present in the majority of top candidate models (Table 1).

**Figure 3.**
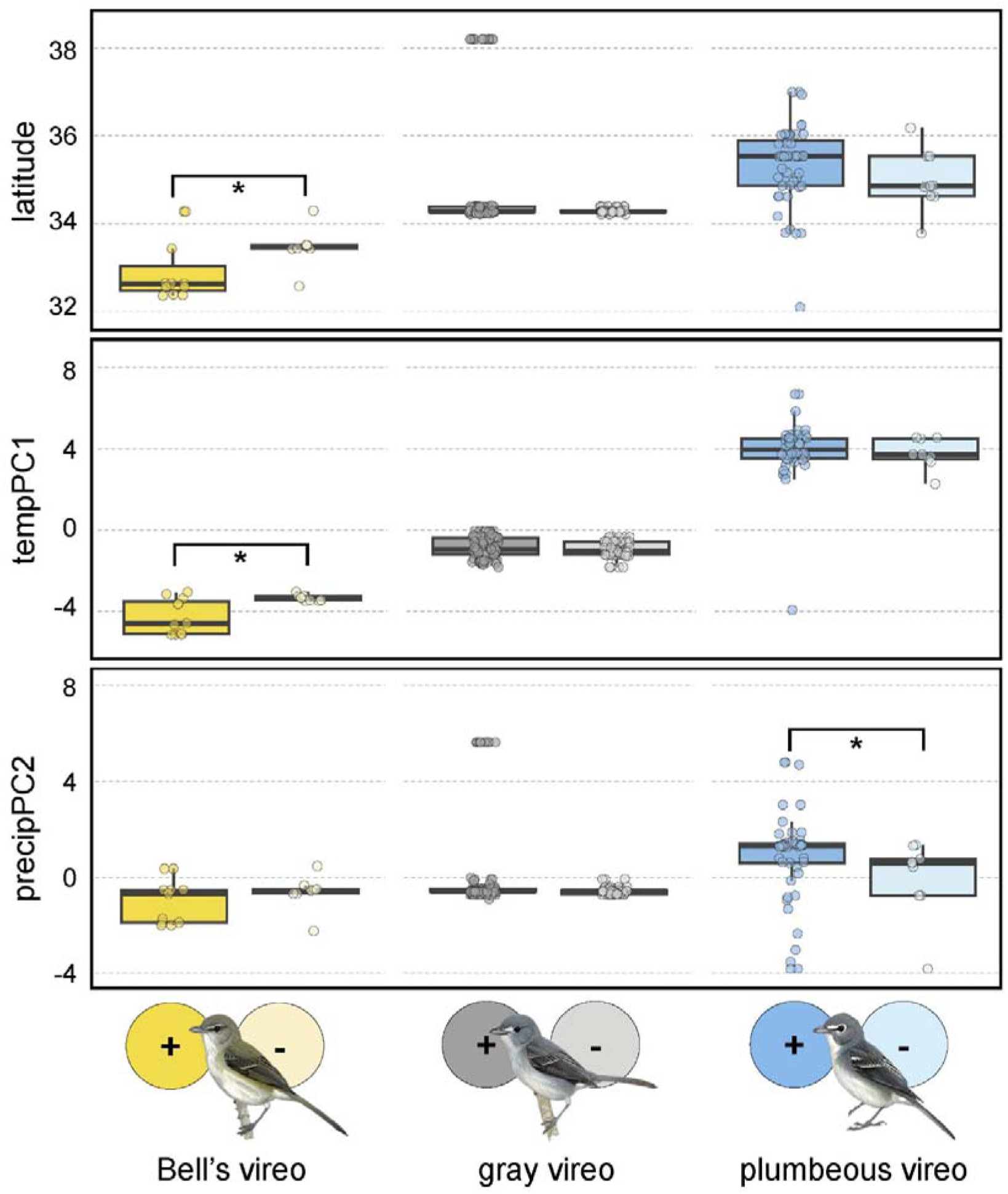
Intraspecific distribution of infected and uninfected vireos across associated environmental variables. Higher tempPC1 values relate to cooler overall temperatures and higher precipPC2 relates to sites with more seasonally stable precipitation. Significantly different distributions are denoted by an asterisk (p < 0.05).

We did not find any association between parasitemia and environmental variables in plumbeous vireos, with the null model having the strongest support and no clear predictor variable significantly reported across top models (Table S10). gray vireos, however, showed a significant pattern of increased parasitemia as overall precipitation (precipPC1) increased (Table S10), but this model had low explanatory power (R² = 0.05).

## Discussion

Vireo species breeding in the southwestern U.S. are especially susceptible to haemosporidian infection, particularly by parasites in the *Parahaemoproteus* subgenus. In this study, we uncovered that the elevational limits of haemosporidian parasites closely parallel those of their avian hosts. We observed striking differences in host specificity and parasite richness across host species and their associated habitats, revealing that the high-elevation plumbeous vireos harbored the most diverse and phylogenetically distinct parasite lineages, especially compared to another well-sampled, lower elevation host species, the gray vireo. Although parasite prevalence differed significantly between Bell’s vireos (55% infected) and plumbeous vireos (86.2% infected), which breed under different levels of precipitation and temperature, we found variable links between infection status and climatic factors in our dataset. Specifically, we found that infection probability was strongly influenced by aspects of temperature for Bell’s vireos, elevation for gray vireos, and precipitation for plumbeous vireos, likely reflecting subtle differences in habitat suitability for arthropod vectors. Lastly, we found low to moderate parasitemia values in both gray and plumbeous vireos, consistent with chronic infections. We examine these results in relation to previous haemosporidian research in the southwest U.S. and in vireos, both in their breeding and nonbreeding grounds, and discuss remaining questions in the sections below.

### Elevational replacement and haemosporidian diversity patterns

Avian haemosporidians exhibit variable patterns of turnover across elevations (Galen and Witt, 2014; Illera et al., 2015; McNew et al., 2021), which is often independent of geographic distance between hosts (Williamson et al., 2019; Barrow et al., 2021). We demonstrated a strong case of parallel elevational replacement in hosts and parasites, with high turnover among *Parahaemoproteus* assemblages infecting vireo species breeding at different elevations. This elevational replacement pattern could be the result of a few different scenarios. For example, shared historical and ecological forces, such as habitat specialization in both hosts and vectors (Ellis et al., 2020), may shape the elevational range limits of both vireos and their *Parahaemoproteus* parasites. Alternatively, species-specific factors that co-evolved independently in response to specialized parasite lineage pressures, such as immune defenses (e.g., Armour et al., 2025), could also be at play. Whether parasite elevational replacement is host-driven or independently mediated by the environmental gradient, global patterns of spatial diversity in parasites should be tightly coupled with those of their hosts.

Of the 30 cyt *b Parahaemoproteus* and 12 *Plasmodium* haplotypes currently reported to infect vireo species (MalAvi; downloaded 1/1/2025), our study adds seven novel *Parahaemoproteus* and one novel *Plasmodium* haplotype. The near absence of *Plasmodium* infections, even at lower elevations, contrasts findings from other temperate region studies (e.g. in Alaska: Oakgrove et al., 2014; California: Walther et al., 2016; Oklahoma: Wyckoff et al., 2024) but is consistent with previous screening at high elevations in New Mexico (Marroquin-Flores et al., 2017; Williamson et al., 2019; Barrow et al., 2021). While the cooler temperatures experienced at higher elevations have been proposed as a potential limiting factor for haemosporidian infection (e.g. LaPointe et al., 2012; Barrow et al., 2021), the low prevalence of *Plasmodium* at lower elevations in this region may also reflect limited suitable habitat for mosquito vectors prior to the onset of monsoon rains. Broader and temporally replicated surveys of potential vectors and vector abundance in this region are needed to fully explain this apparent absence.

We identified an interesting pattern among the *Parahaemoproteus* lineages found in our focal species: plumbeous vireos harbored a broad diversity of parasite haplotypes relative to the other two sampled species, while gray vireos exhibited a depauperate and likely specialized haemosporidian assemblage (Fig. 1D). This disparity suggests that plumbeous vireos may support more permissive or ecologically flexible parasite associations, while gray vireos may be undergoing, or may have already undergone, a transition toward fixed symbiosis with a narrower set of haemosporidian lineages. The contrasting pattern of haemosporidian diversity between vireo species could alternatively be caused by differences in avian host richness, and the low diversity found in gray vireos could reflect the depauperate breeding bird communities of the arid and structurally simple juniper savanna habitats.

### Conserved high susceptibility to haemosporidians among vireo species

Vireos stand out among southwestern songbirds for their consistently high haemosporidian infection rates, providing a valuable opportunity to examine parasite dynamics in hyper-susceptible hosts. In the three species screened here, we found that 70.6% of 248 individuals screened had at least one haemosporidian infection, which is greater than double the prevalence reported in broader community-wide surveys conducted in the southwestern U.S. over a similar period. For example, Marroquin-Flores et al. (2017) screened 186 birds of 49 species sampled in New Mexico from June-July 2016, including 13 individual vireos of two species: *n* = 4 warbling vireos (*Vireo gilvus*) and *n* = 9 plumbeous vireos. The average proportion of infection was 36.6% across all studied species, but 92% of vireos tested positive. A subsequent large-scale study by Barrow et al. (2021) reported similar patterns, with an overall haemosporidian prevalence of 36.1% among 776 total individuals of 61 species. Of these, vireos again exhibited significantly higher infection proportions, with plumbeous and warbling vireos demonstrating the highest prevalence in the study at 87% and 78%, respectively (Barrow et al., 2021).

This observation of high susceptibility in vireos is not limited to birds sampled in the southwestern U.S., as other studies across breeding sites have also reported higher relative prevalences in the *Vireo* genus. Granthon and Williams (2017) reported that 100% of the 62 PCR-screened red-eyed vireos (*Vireo olivaceus*) from Pennsylvania were infected with *Plasmodium*, *Haemoproteus*, or *Leucocytozoon* species (Granthon and Williams, 2017). Rodriquez et al. (2021) found *Haemoproteus* prevalence of 59% in 22 individual warbling vireos sampled from Colorado breeding sites (Rodriguez et al., 2021). Shared susceptibility within closely related species has been extensively documented (e.g. González et al., 2014; Lutz et al., 2015; Barrow et al., 2019; Pigeault et al., 2022) but attempts to identify the specific factors that contribute to high prevalence in vireos are still lacking. Of the conserved host traits recorded to have consistent interaction with haemosporidian prevalence in other avian clades, thus far, shared life history traits such as open cup nest type and insectivorous diets (as reported and reviewed in Braga et al., 2011; Rodriguez et al., 2021) may increase exposure risk, at least partially explaining the high prevalence in this group.

However, a combination of traits, including behavior (e.g., time spent incubating and/or brooding at the nest), features of nest microhabitat (e.g., height of nest from ground), and aspects of immunity (e.g., haemosporidian tolerance) may also contribute to the elevated susceptibility (González et al., 2014; Lutz et al., 2015; Muriel, 2020; Rodriguez et al., 2021; Pigeault et al., 2022).

Interestingly, of the 14 gray vireos that we recaptured and screened for parasites across two sampling years (2017 and 2018), 12 were positive via PCR in both years, and the remaining two birds became infected by the second year (Table S2). Although more data is needed to determine whether these represent chronic infections or reinfections, and pinpoint where transmission occurs, these results indicate gray vireos likely experience sustained parasite pressure during the breeding season. Continued monitoring of haemosporidian infection across vireo populations, coupled with investigations into the mechanisms underlying their susceptibility, could yield valuable insights into the spatial and temporal dynamics of infection risk in this clade.

### Environmental determinants of haemosporidian infection vary by host species

Previous studies have found strong links between haemosporidian prevalence and topographic, geographic, and climatic variables (Pérez-Rodríguez et al., 2013; Rodriguez et al., 2021), but the direction and magnitude of these effects vary by region and haemosporidian clade (Williamson et al., 2019). In the arid southwest, we expected haemosporidian prevalence to correspond with water availability, which is important for insect vector (Valkiūnas, 2004; Santiago-Alarcon et al., 2012), such as those associated with potential water sources (i.e. riparian habitats) and areas with higher, more seasonally stable precipitation (i.e. montane woodlands).

Somewhat contrary to expectations given predictions about the relative aridity of habitat during the pre-monsoon sampling period, we found that interspecific differences in prevalence increased linearly with elevation and precipitation: Bell’s vireos showed the lowest prevalence and gray vireos exhibited unexpectedly high infection rates. We were unable to determine whether observed infection patterns were driven by species-specific traits or abiotic environmental differences, and increased sampling of Bell’s vireos may help clarify or reduce these differences. Within-species analyses revealed no consistent predictors of infection across all taxa (Fig. S2-6); instead, different environmental variables were associated with infection in each species (Fig. 3; Table S9-S10).

Expanded screening of haemosporidians across low- to mid-elevation habitats, among co-distributed species, at variable distance from water sources, and across precipitation gradients would further elucidate cross-elevational trends and enhance predictions of infection probability in avian communities of the southwestern U.S.

### Moderate and consistent infection loads across environmental gradients

Gray and plumbeous vireos exhibited similar average infection loads (∼1% in gray vireos, 0.786% in plumbeous, range: 0.03–4.5%) with no significant differences between species (Fig. 1B). These values slightly exceed the regional cross-species average (<1%; Marroquin-Flores et al., 2017) and fall within the range of moderate infection intensities reported in experimental studies of other passerines (e.g. Asghar et al., 2012). We did note, however, two gray vireos with elevated parasitemia (4.5 and 4.49%) were sampled at distinct sites in the Sevilleta National Wildlife Refuge. The elevated parasitemia in these individuals likely represented acute infection acquired shortly before capture and prior to immune response, or recrudescence of chronic infections exacerbated by physiological stressors such as nutritional deficits or hormonal fluctuations (Arriero et al., 2018; Names et al., 2021). The absence of high parasitemia in Bell’s and plumbeous vireos could be an artifact of limited sampling, as birds with severe infections often exhibit reduced mobility and capture probability (Mukhin et al., 2016), suggesting that heavily infected individuals in these species may be underrepresented.

Additionally, we found no relationship between parasitemia and any tested environmental variables for plumbeous vireos. In gray vireos we saw a significant linear trend in parasitemia with increased overall precipitation (precipPC1; Table S10), but this model had weak explanatory power (R² = 0.05). These results are consistent with previous findings which suggest that haemosporidian parasitemia patterns are shaped by complex interactions between host and parasite biology not easily captured in models with abiotic factors alone (Loiseau et al., 2013).

### Conclusions

We found that parasite elevational distributions mirror the elevational replacement pattern of host species among vireos in southwestern North America. This pattern may have resulted from host specialization, climate variation independently acting on hosts and parasites, climate variation acting indirectly through habitat or broader host and vector communities, or a combination of those forces. We found that vireos were consistently infected at high rates and largely harbor chronic, low-to-moderate intensity infections dominated by *Parahaemoproteus*, suggesting the potential for exclusive, reciprocal selective pressures that facilitate host-parasite coadaptation. The contrast in parasite richness between plumbeous vireos and gray vireos was striking: gray vireos harbored a limited number of haemosporidian lineages, all of which were host specific, while plumbeous vireos were infected with a diverse set of lineages. These differences imply species-specific variability in resistance, tolerance, or exposure to generalist haemosporidian lineages. In sum, our work adds to a growing body of evidence about clade-specific avian host susceptibility and host-parasite turnover across elevational gradients, highlighting vireos as an ideal focal group for future work on host-parasite dynamics across space and time.

## Acknowledgements

The authors sincerely thank Bethany Abrahamson, Michael Andersen, Matthew Baumann, Alison Boyer, Serina Brady, Sara Brant, Mariel Campbell, Andrea Chavez, Mina Carnicom, Lida Crooks, Rosario Marroquin-Flores, Chauncey Gadek, Spencer Galen, Ethan Gyllenhaal, Mike Hartshorne, Andy Johnson, Michael Lelevier, Joseph Manthey, Xena Mapel, Taylor Martinez, Jenna McCullough, Moses Michelsohn, Kathleen Ramsay, George Rosenberg, C. Jonathan Schmitt, Donna Schmitt, Brenda Villanueva, and Toby Weisenhaus.

## Declarations

Not applicable.

## Ethics approval

Research activities were conducted under Institutional Animal Care and Use Protocol 16-200406-MC and appropriate state and federal scientific collecting permits (New Mexico Department of Game and Fish Authorization Number 3217; US Fish and Wildlife Permit Numbers MB094297-0 and MB0942970-1).

## Availability of data and material

Specimen data are available in Supplementary Tables S1–S10 and are searchable via Arctos (arctos.database.museum; Cicero et al., 2024). Parasite sequences are archived in GenBank (accession nos. PV295634-PV295830). Vouchered specimens and frozen tissue samples are archived at the Museum of Southwestern Biology Division of Birds and Division of Genomic Resources, respectively (https://msb.unm.edu/). All R code and datasets are available on FigShare (https://doi.org/10.6084/m9.figshare.28585943).

## Authors’ contributions

DLFW, JLW, CCW, and LNB conceptualized and developed methodology for this project. JLW, SEF, HMS, SMB, KG, CCW, and LNB performed fieldwork and subsequent data curation. DLFW, JLW, and SMB performed molecular analysis and generated sequence data. JLW, SEF, HMS, KG, CCW, and LNB secured funding and provided supervision. DLFW and JLW performed formal analyses and created visualizations. DLFW and JLW wrote the original draft of the manuscript, and DLFW wrote the final version with input from all co-authors.

