## Supplemental Figures 1-6 for "Parallel elevational replacement of hosts and parasites in a highly susceptible songbird genus" (from another version of this preprint: version 2, 2025-10-16)

**LIST OF SUPPLEMENTAL FIGURES**

Figure S1: Principal Components Analysis of Bioclimatic variables.

Figure S2: Distribution of infected and uninfected across latitudes for all Vireos, Bell’s, Gray, and Plumbeous Vireos.

Figure S3: Distribution of infected and uninfected across tempPC1 for all Vireos, Bell’s, Gray, and Plumbeous Vireos.

Figure S4: Distribution of infected and uninfected across tempPC2 for all Vireos, Bell’s, Gray, and Plumbeous Vireos.

Figure S5: Distribution of infected and uninfected across precipPC1 for all Vireos, Bell’s, Gray, and Plumbeous Vireos.

Figure S6: Distribution of infected and uninfected across precipPC2 for all Vireos, Bell’s, Gray, and Plumbeous Vireos.

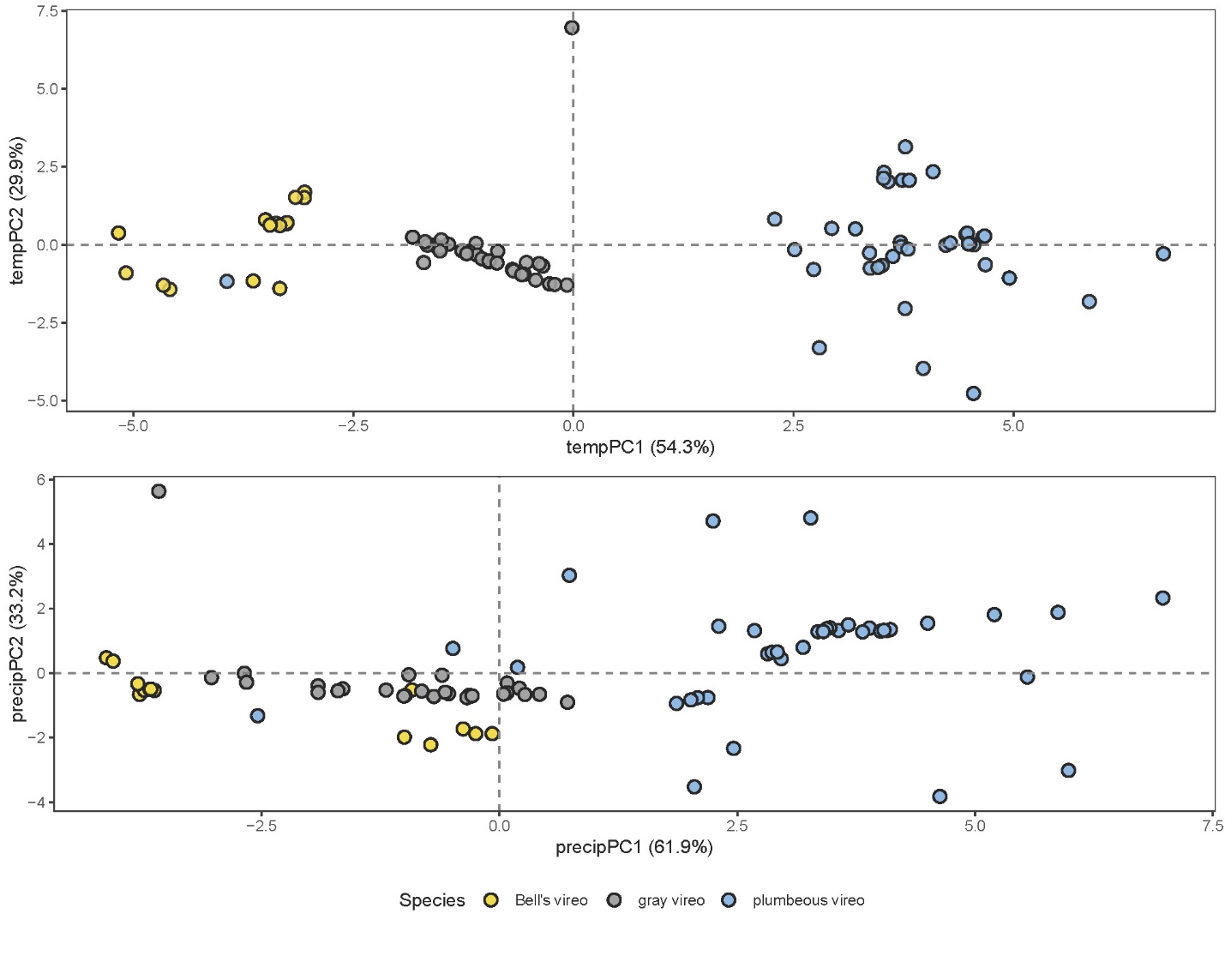

**Figure S1.** First two Principal Components Analysis axes of temperature (Bio 1-10; upper, tempPC) and precipitation (Bio11-l9; lower, precipPC), colored by species (WorldClim 2.1; Fick & Hijmans, 2017). TempPC1 primarily represents year-round temperatures, with higher values relating to lower averages, while tempPC2 represents greater seasonality with higher values relating to more extreme fluctuations throughout the year and day-night cycle. PrecipPC1 largely represents overall precipitation, with higher values indicating wetter conditions, whereas high precipPC2 corresponds with more stable and less seasonal precipitation. The outlier Gray Vireo points in both temperature and precipitation represent birds sampled from the Abajo Mountains site in Utah.

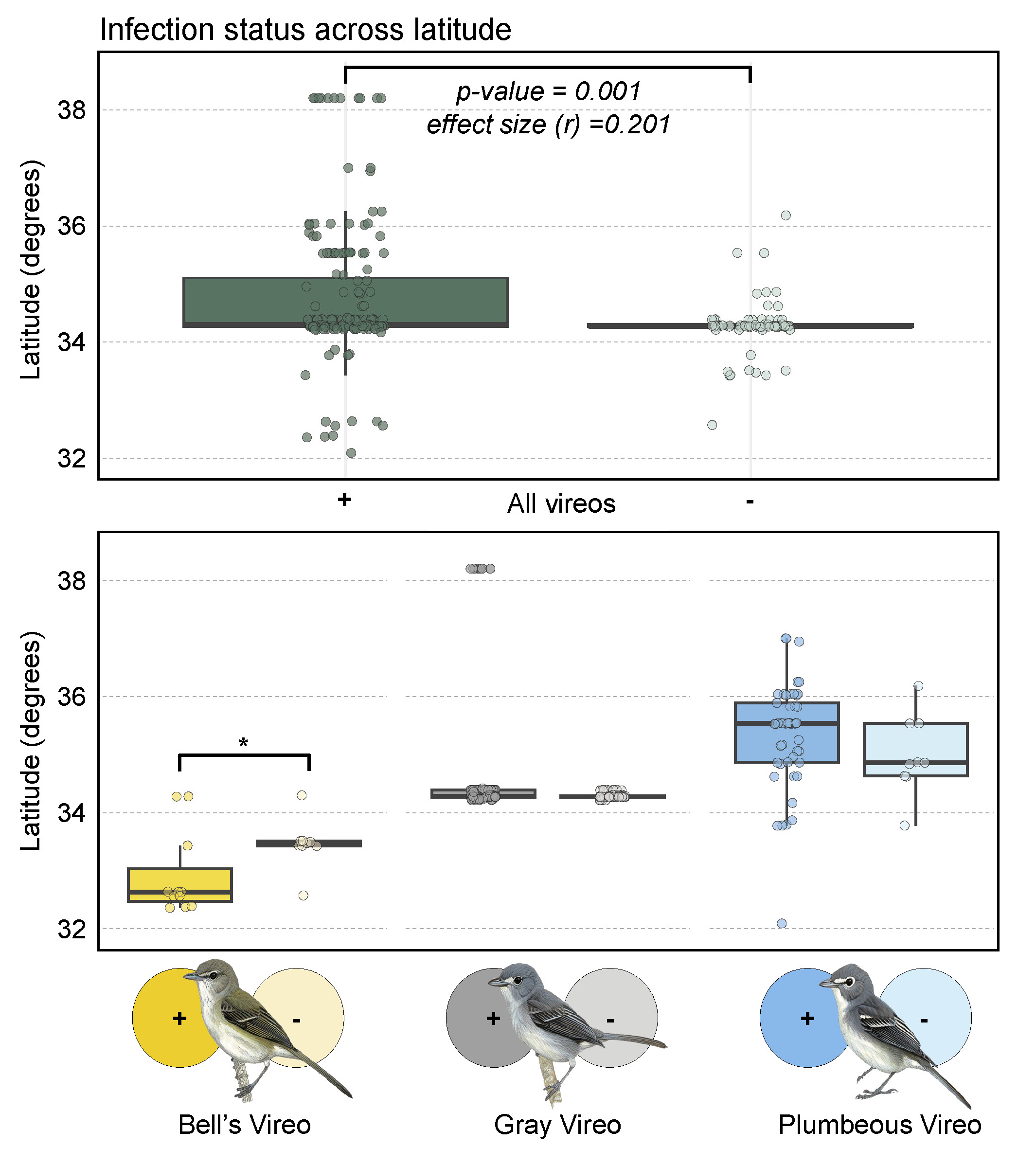

**Figure S2.** Distribution of infected (darker shade) and uninfected (lighter shade) across latitudes for all vireos, Bell’s, Gray, and Plumbeous. Mann-Whitney U-test p-values are shown where significant (p < 0.05). Wilcoxon effect size shown for all species comparison.

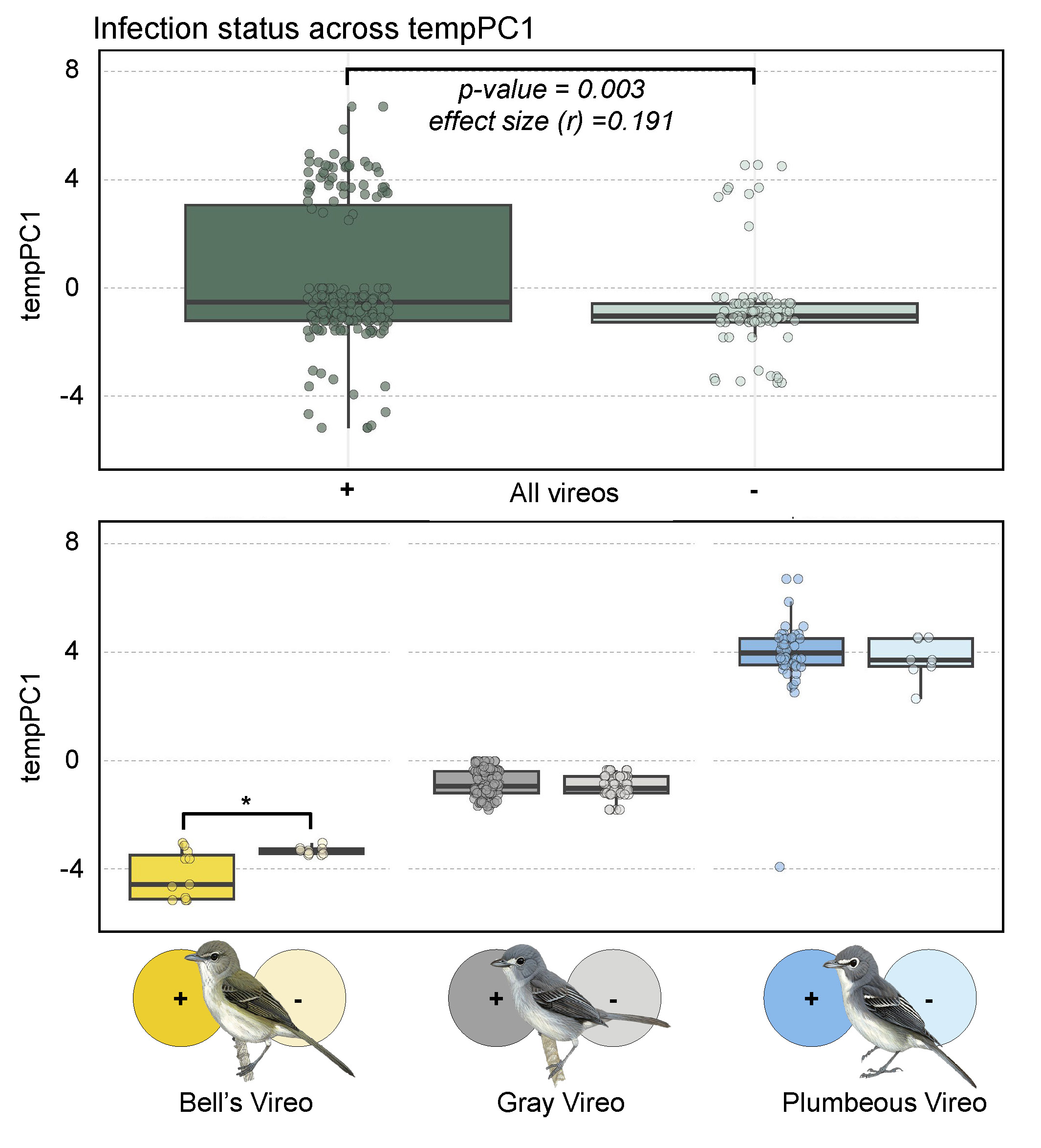

**Figure S3.** Distribution of infected (darker shade) and uninfected (lighter shade) across tempPC1 for all vireos, Bell’s, Gray, and Plumbeous. Higher tempPC1 values relating to lower averages. Mann-Whitney U-test p-values are shown where significant (p < 0.05). Wilcoxon effect size shown for all species comparison.

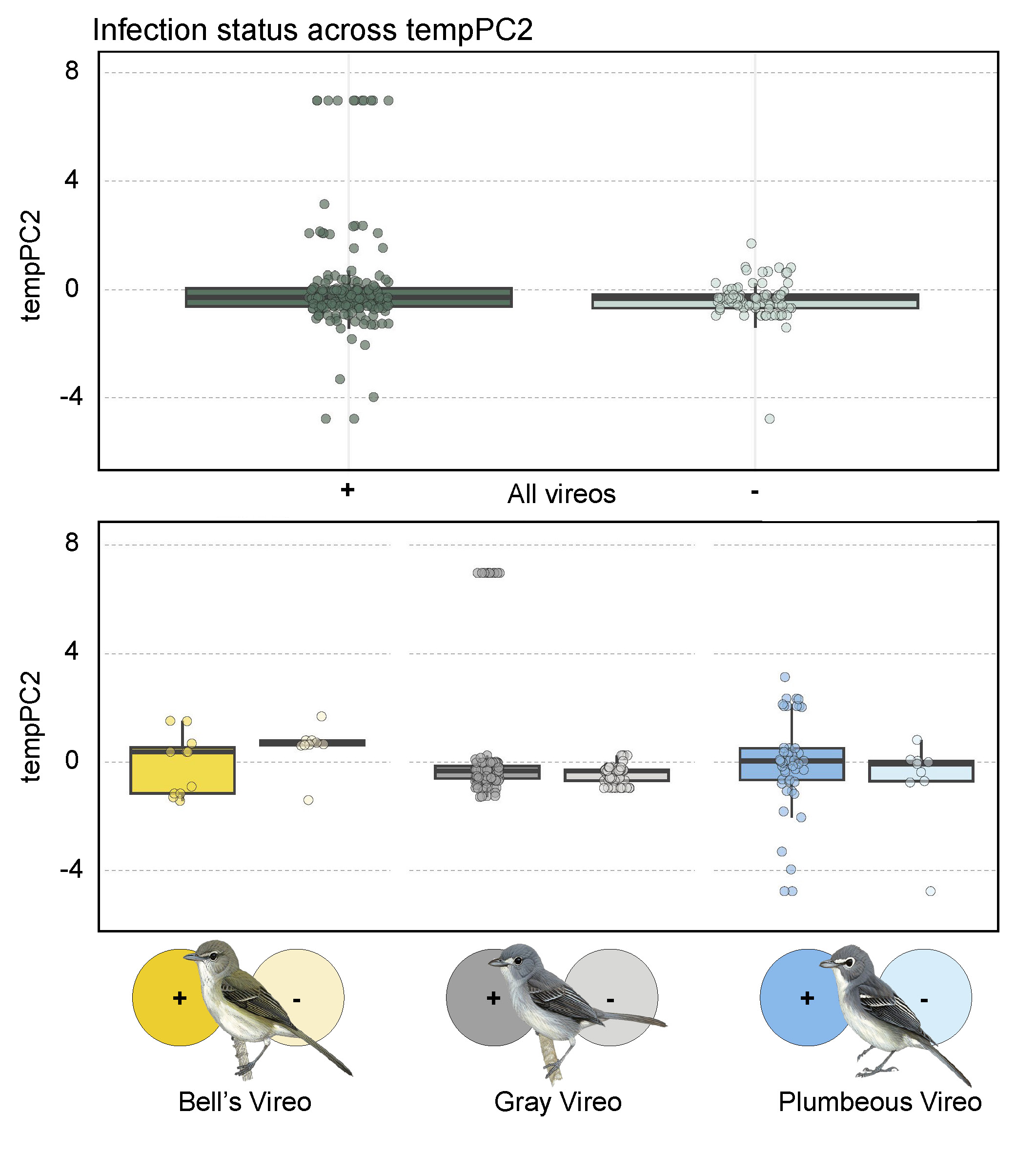

**Figure S4.** Distribution of infected (darker shade) and uninfected (lighter shade) across tempPC2 for all vireos, Bell’s, Gray, and Plumbeous. Higher tempPC2 values relate to more extreme temperature fluctuations throughout the year and day-night cycle. Mann-Whitney U-test p-values are shown where significant (p < 0.05).

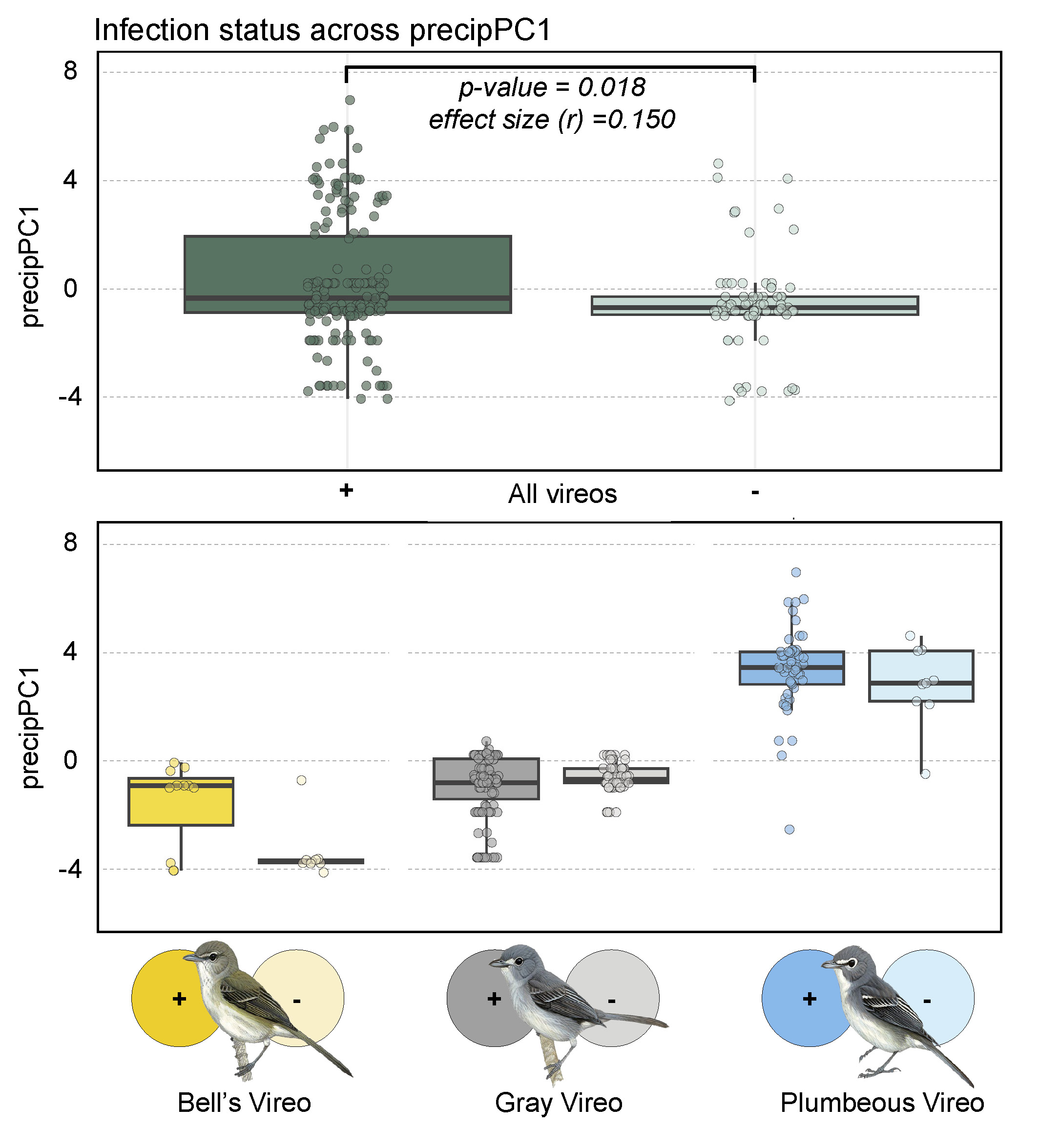

**Figure S5.** Distribution of infected (darker shade) and uninfected (lighter shade) across tempPC2 for all vireos, Bell’s, Gray, and Plumbeous. Higher precipPC1 values relate to wetter conditions, overall. Mann-Whitney U-test p-values are shown where significant (p < 0.05). Wilcoxon effect size shown for all species comparison.

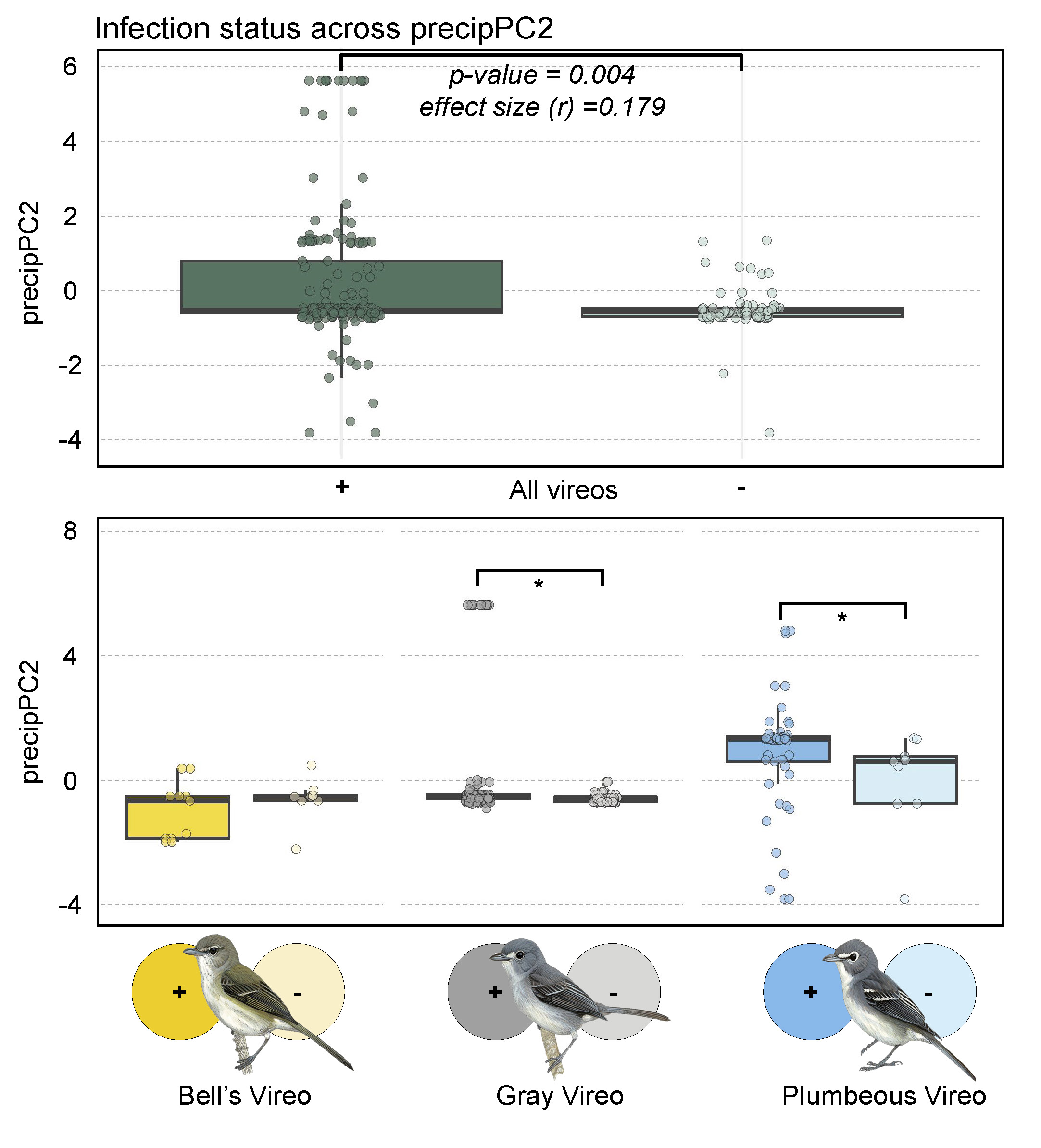

**Figure S6.** Distribution of infected (darker shade) and uninfected (lighter shade) across tempPC2 for all Vireos, Bell’s, Gray, and Plumbeous. Higher precipPC2 corresponds with more stable and less seasonal precipitation. Mann-Whitney U-test p-values are shown where significant (p < 0.05). Wilcoxon effect size shown for all species comparison.
