## Supplemental Tables 1-11 for "Parallel elevational replacement of hosts and parasites in a highly susceptible songbird genus" (from another version of this preprint: version 2, 2025-10-16)

**LIST OF SUPPLEMENTAL TABLES**

Table S1: Sample information for all host individuals and parasite cytochrome b haplotypes (separate excel spreadsheet).

Table S2: Record of Gray Vireo recapture, including parasite infection status and parasitemia (separate excel spreadsheet).

Table S3: Documentation of manual coordinate assignments for subset of samples not assigned coordinates at the time of collection.

Table S4: Haemosporidian haplotype diversity and abundance by host, cytochrome b consensus sequences for haplotypes, and number of other host species infected by each haplotype found on MalAvi as of 3/1/2025 (separate excel spreadsheet).

Table S5: PCA loadings and interpretations relating to temperature.

Table S6: PCA loadings and interpretations relating to precipitation.

Table S7: Parasite richness output from iNEXT R-script.

Table S8: Sampling completeness output from iNEXT R-script.

Table S9: Summary of statistical analyses used to compare infection status across six variables.

Table S10: Summary of generalized linear model (GLM) results assessing the relationship between environmental variables and haemosporidian infection parasitemia in Gray and Plumbeous Vireos.

Table S11: Cytochrome b sequence base pair percentage differences, measured as Jukes-Cantor distances, between haemosporidian haplotypes (separate excel spreadsheet).

**Supplemental Table S2.** Record of Gray Vireo recapture for 14 individuals, including parasite infection status and parasitemia. Birds were initially captured in 2017 and recaptured in 2018. Rows with the “PCR_pos” column value in red represent samples that were sent for sequencing and catalogued in GenBank. See separate Excel spreadsheet.

| **Band number** | **Sampling locality** | **Assigned latitude** | **Assigned longitude** |
| --- | --- | --- | --- |
| 272132433 | Cottonwood | 34.2686 | -106.6336 |
| 272132447 | Montoso | 34.2686 | -106.6336 |
| 272132629 | Montoso | 34.2686 | -106.6336 |
| 272132640 | Montoso | 34.2686 | -106.6336 |
| 272132641 | Montoso | 34.2686 | -106.6336 |
| 272132636 | Pinyon Canyon | 34.38896 | -106.5546 |
| 272132637 | Pinyon Canyon | 34.38896 | -106.5546 |
| 272132652 | Pinyon Canyon | 34.38896 | -106.5546 |
| 272132653 | Pinyon Canyon | 34.38896 | -106.5546 |
| 272132654 | Pinyon Canyon | 34.38896 | -106.5546 |
| 272132665 | Pinyon Canyon | 34.38896 | -106.5546 |
| 272132679 | Pinyon Canyon | 34.38896 | -106.5546 |
| 272132656 | Private Montoso | 34.2982 | -106.577 |
| 272132661 | Private Montoso | 34.2982 | -106.577 |
| 272132412 | Sepultura Flats | 34.2686 | -106.6336 |
| 272132415 | Sepultura Flats | 34.2686 | -106.6336 |
| 272132416 | Sepultura Flats | 34.2686 | -106.6336 |
| 272132417 | Sepultura Flats | 34.28732 | -106.61492 |
| 272132626 | Sepultura Flats | 34.2686 | -106.6336 |
| 272132634 | Sepultura Flats | 34.2686 | -106.6336 |
| 272132434 | Tomasino | 34.2313 | -106.6752 |
| 272132630 | Tomasino | 34.2313 | -106.6752 |
| 272132639 | Tomasino | 34.2313 | -106.6752 |
| 272132642 | Tomasino | 34.2313 | -106.6752 |
| 272132643 | Tomasino | 34.2313 | -106.6752 |
| 272132644 | Tomasino | 34.2313 | -106.6752 |
| 272132647 | Tomasino | 34.2313 | -106.6752 |
| 272132650 | West Side | 34.3917 | -107.0095 |

**Supplemental Table S3. Manual assignment of coordinates for Gray Vireos without sample locality at the time of collection**. Note: All Gray Vireos that were missing coordinates were collected <100m from the closest sampling locality. Therefore, for birds missing coordinates from the Sepultura Flats area, we used intermediate coordinates between Sepultura, Montoso, and Cottonwood: 34.2686, -106.6336. For Pinyon Canyon birds missing coords, we used the following: 34.38896, -106.5546, Private Montoso sampling were located ~1.87 km apart. For two birds with missing Private Montoso coordinates, we selected the midpoint between the two coords: 34.2982, -106.577. For Tomasino birds missing coords, we used the following: 34.2313, -106.6752. Single missing West Side bird given coordinates intermediate to the other two: 34.3917, -107.0095.

**Supplemental Table S4.** Haemosporidian haplotype diversity and abundance by host, cytochrome b consensus sequences for haplotypes, and number of other host species infected by each haplotype found on MalAvi as of 3/1/2025. See separate Excel spreadsheet.

| **Temperature PCA Loadings** | | |
| --- | --- | --- |
| **Dimension 1 (PC1)** | **53.9% variation captured** | |
| *Contributing variables* | *Loadings* | *Interpretation as TPC1 increases* |
| BIO1: Annual Mean Temp | -0.406 | Colder annual temps |
| BIO8: Mean Temp of Wettest Quarter | -0.400 | Colder wet quarter |
| BIO11: Temp of Coldest Quarter | -0.396 | Colder coldest quarter |
| BIO10: Mean Temp of Warmest Quarter | -0.385 | Colder warm quarter |
| BIO6: Min Temp of Coldest Month | -0.373 | Colder cold month |
| BIO5: Max Temp of Warmest Month | -0.365 | Colder warm month |
| **Dimension 2 (PC2)** | **28.4% variation further captured** | |
| *Contributing variables* | *Loadings* | *Interpretation as TPC2 increases* |
| BIO4: Temp Seasonality | 0.549 | More seasonal temps |
| BIO7: Temp Annual Range | 0.529 | Greater range of annual temps |
| BIO9: Mean Temp of Driest Quarter | 0.371 | Warmer temps during driest quarter |
| BIO3: Isothermality | -0.370 | Greater difference in day-night temps |

**Supplemental Table S5. First and second dimension PCA loadings and interpretations relating to temperature (tempPC1-2).** Variables, loadings, and interpretation of temperature PC axes that contribute above the average contribution from all variables (i.e., 1/11 or 9.09%) listed in descending order. Loadings show the relative contribution and direction (positive or negative value) of the variable relationship to an increasing PC axis.

| **Precipitation PCA Loadings** | | |
| --- | --- | --- |
| **Dimension 1 (PC1)** | **61.9% variation captured** | |
| *Contributing variables* | *Loadings* | *Interpretation as PPC1 increases* |
| BIO12: Annual Precip | 0.444 | More annual precip |
| BIO14: Precip of Driest Month | 0.391 | More precip in driest month |
| BIO17: Precip of Driest Quarter | 0.365 | More precip in driest quarter |
| BIO18: Precip of Warmest Quarter | 0.361 | More precip in warmest quarter |
| BIO19: Precip of Coldest Quarter | 0.357 | More precip in coldest quarter |
| BIO13: Precip of Wettest Month | 0.357 | More precip in wettest month |
| **Dimension 2 (PC2)** | **33.2% variation further captured** | |
| *Contributing variables* | *Loadings* | *Interpretation as PPC2 increases* |
| BIO15: Precip Seasonality | -0.582 | Not as seasonal of precip |
| BIO16: Precip of Wettest Quarter | -0.397 | Less precip in wettest quarter |
| BIO13: Precip of Wettest Month | -0.367 | Less precip in wettest month |
| BIO18: Precip of Warmest Quarter | -0.357 | Less precip during warmest quarter |

**Supplemental Table S6. First and second dimension PCA loadings and interpretations relating to precipitation (precipPC1-2).** Variables, loadings, and interpretation of precipitation PC axes that contribute above the average contribution from all variables (i.e., 1/8 or 12.5%) listed in descending order. Loadings show the relative contribution and direction (positive or negative value) of the variable relationship to an increasing PC axis.

| **Assemblage** | **Diversity** | **Observed** | **Estimator** | **s.e.** | **LCL** | **UCL** |
| --- | --- | --- | --- | --- | --- | --- |
| Bell's Vireo | species richness | 3 | 3.92 | 0.784 | 3.00 | 5.45 |
| Gray Vireo | species richness | 4 | 4.99 | 0.438 | 4.14 | 5.84 |
| Plumbeous Vireo | species richness | 14 | 53.8 | 23.3 | 14.0 | 101 |

**Supplemental Table S7.** **iNEXT estimates of haemosporidian parasite species richness for each vireo species.** The table includes observed and estimated species richness, standard error, and 95% confidence intervals.

| **Assemblage** | **N host** | **SC** | **m** | **Method** | **qD** | **qD.LCL** | **qD.UCL** |
| --- | --- | --- | --- | --- | --- | --- | --- |
| Bell's Vireo | 12 | 0.996 | 33 | Extrapolation | 3.89 | 1.83 | 5.95 |
| Gray Vireo | 123 | 0.957 | 14 | Rarefaction | 1.81 | 1.15 | 2.47 |
| Plumbeous Vireo | 62 | 0.952 | 364 | Extrapolation | 40.6 | 10.9 | 70.2 |

**Supplemental Table S8.** **iNEXT summary of haemosporidian parasite diversity estimates at approximately 95% sample completeness for each vireo species.** The table includes the number of infections sampled, estimated sample completeness (SC), the number of infections required to reach 95% sampling completeness (m), the estimation method used, and diversity estimates with 95% confidence intervals.

| **Infected vs. uninfected** | | | | | | | | | | | | |
| --- | --- | --- | --- | --- | --- | --- | --- | --- | --- | --- | --- | --- |
|  | Bell's Vireo | | | | Gray Vireo | | | | Plumbeous Vireo | | | |
|  | *p* | *W* | *r* | *z* | *p* | *W* | *r* | *z* | *p* | *W* | *r* | *z* |
| *Latitude* | **0.040** | 77 | 0.467 | -2.09 | 0.086 | 2647 | - | 1.72 | 0.119 | 2775 | - | 1.57 |
| *Elevation* | 0.148 | 30 | - | -1.48 | **0.038** | 2541 | 0.159 | -2.07 | 0.772 | 207 | - | -0.301 |
| *tempPC1* | **0.027** | 79 | 0.502 | -2.25 | 0.196 | 2775 | - | 1.29 | 0.396 | 181 | - | 0.859 |
| *tempPC2* | 0.110 | 71 | - | -1.64 | 0.21 | 2787 | - | 1.25 | 0.308 | 173 | - | 1.03 |
| *precipPC1* | 0.093 | 27 | - | 1.72 | 0.578 | 3325 | - | -0.545 | 0.339 | 176 | - | 0.967 |
| *precipPC2* | 0.402 | 61 | - | -0.877 | 0.057 | 2597 | - | 1.90 | **0.042** | 126 | 0.268 | 2.04 |

**Supplemental Table S9. Summary of statistical analyses used to compare infection status across six variables.** Mann-Whitney U-test statistics (W), z-value (z), p-values (p), and effect sizes (r) for analyses of infection status (infected vs. uninfected) within Bell's Vireo (*Vireo bellii*, *n* = 20), Gray Vireo (*V. vicinior*, *n* = 170), and Plumbeous Vireo (*V. plumbeus*, *n* = 58). Bold text indicates significance (p < 0.05).

| ***Top five parasitemia fixed effects additive models*** | | | | | |
| --- | --- | --- | --- | --- | --- |
| **Gray Vireo (*n* = 51)**  Best model: higher parasitemia as precipitation increases; R² = 0.05 | | | | | |
| *Rank* | *Predictors* | *K* | *AICc* | *ΔAICc* | *Weight* |
| *1* | precipPC1 | 3 | 21.1 | *-* | 0.224 |
| *2* | precipPC1, tempPC1 | 4 | 21.1 | *0.044* | 0.219 |
| *3* | tempPC2 | 3 | 21.4 | *0.254* | 0.197 |
| *4* | Elevation | 3 | 21.5 | *0.380* | 0.185 |
| *5* | precipPC2 | 3 | 21.6 | *0.511* | 0.174 |
| **Plumbeous Vireo *n* = 28**  Best model: Intercept only (null) model | | | | | |
| *Rank* | *Predictors* | *K* | *AICc* | *ΔAICc* | *Weight* |
| *1* | Null model | 2 | 16.3 | *-* | 0.267 |
| *2* | precipPC2 | 3 | 17.9 | *1.57* | 0.122 |
| *3* | tempPC2 | 3 | 18.0 | *1.74* | 0.112 |
| *4* | precipPC1 | 3 | 18.3 | *1.94* | 0.101 |
| *5* | Elevation | 3 | 18.7 | *2.37* | 0.082 |

**Supplemental Table S10. Top five models from information-theoretic model selection assessing environmental predictors of haemosporidian parasitemia in Gray Vireos (*Vireo vicinior*).** Models are ranked by AICc, with ΔAICc values relative to the top model and corresponding Akaike weights (Weight) indicating model support.

**Supplemental Table S11:** Cytochrome b sequence base pair percentage differences, measured as Jukes-Cantor distances, between haemosporidian haplotypes. In separate Excel spreadsheet.
